# Contributions of hippocampus and striatum to memory-guided behavior depend on past experience

**DOI:** 10.1101/041459

**Authors:** Janina Ferbinteanu

## Abstract

The hippocampal and striatal memory systems operate independently and in parallel in supporting spatial and response learning, respectively, when animals are explicitly trained in one task. Here, we investigated whether this principle continues to hold when animals are concurrently trained in two types of tasks. Rats were trained on a plus maze in either a spatial navigation or a cue response task (individual training), while a third set of rats acquired both (concurrent training). Subsequently, the rats underwent either sham surgery or neurotoxic lesions of the hippocampus (HPC), medial dorsal striatum (DSM) or lateral dorsal striatum (DSL), followed by retention testing. Finally, rats in the individual training condition also acquired the novel ‘other’ task. When rats learned one task, HPC and DSL selectively supported spatial navigation cue response, respectively. However, when rats learned both tasks, HPC and DSL additionally supported the behavior incongruent with the processing style of the corresponding memory system. DSM significantly contributed to performance regardless of task or training procedure. Experience with the cue response task facilitated subsequent spatial learning, while experience with spatial navigation delayed both simultaneous and subsequent response learning. These findings suggest that multiple principles govern the interactions among memory systems.

**Significance Statement:** Currently, we distinguish among several types of memories, each supported by a distinct neural circuit. The memory systems are thought to operate independently and in parallel. Here, we demonstrate that the hippocampus and the dorsal striatum memory systems indeed operate independently and in parallel when rats learn one type of task at a time, but interact co-operatively and in synergism when rats concurrently learn two types of tasks. Furthermore, new learning is modulated by past experiences. These results can be explained by a model in which independent and parallel information processing that occurs in the separate memory-related neural circuits is supplemented by information transfer between the memory systems at the level of the cortex.

---

Memory is not unitary; rather, there are different types of memories, with different properties and dynamics. The formulation of this organizational principle goes back to the debate between Hull and Thorndike, who proposed that stimulus-response (S-R) associations were sufficient to generate adaptive behavior ^1^ ^2^; and Tolman, who argued that animals additionally form flexible cognitive representations by learning stimulus-stimulus (S-S) associations ^3^. Eventually, Tolman’s view was proved right. It started to become apparent that these memories also involve separate neurobiological substrates when surgical resection of medial temporal lobes in patient H.M. resulted in profound amnesia ^4^ ^5^, a loss of the ability to learn and remember factual information or autobiographical events (declarative memory). Despite the severity of their deficit in the declarative domain, amnesics can form other types of mnemonic representations, as for example memories for motor skills ^6^ ^7^, which are dependent on the neostriatum ^8^, a part of the basal ganglia that is more generally involved in habits ^9^. These and other findings generated the memory systems theory, which posits that different types of memories are supported by distinct neural circuits with individual characteristics of information processing ^10^. Memory systems are considered to operate independently and in parallel to support behavior.

Declarative memories, so named because in humans they can be consciously expressed (declared) verbally, are flexible, form fast, and involve storage of S-S associations. In contrast, habits do not require consciousness, are rigid, form gradually, and are based on remembering S-R associations ^11^ The term ‘declarative’, coined based on research in humans, implies verbal language, but animals also form declarative-like memories. In the rat animal model, spatial learning is representative for declarative memory because it involves a flexible cognitive representation, a spatial cognitive map ^12^. In contrast, habits can be studied by using tasks involving memory for cue-response associations, or memory for body turns. Studying different types of memories by changing the contingencies of the task within the same experimental setting isolates the memory representation guiding behavior by eliminating the effect of confounding variables. Such an approach has been indeed successfully used to demonstrate the operational principle of independent parallelism, and the evidence comes in the form of from double dissociation data, in which damage to brain structure A causes deficits in Task1 but not Task2, while damage to structure B produces the reverse pattern of effects. Much of this work has been directly inspired by Tolman’s experiments using the plus maze ^13^. For example, Packard and McGaugh (1996) used a plus maze whose North arm was blocked and trained rats to walk from the South start arm to find food in the West arm. When placed on the previously blocked North arm early in the training, most rats continued to go to the West arm, demonstrating *place learning*. HPC inactivation rendered the choice random, while DS inactivation had no effect. In contrast, later in training, most rats chose the East arm (they maintained the body turn), demonstrating therefore *response* learning. HPC inactivation had no effect, but DS inactivation resulted in a return to the spatial strategy. Thus, the initially prevalent HPC-dependent spatial strategy was abandoned in favor of a DS-dependent response strategy, suggesting that habits take time to form, but once formed, they dominate behavior.

A similar approach has been used in recording experiments targeting the activity of CA1 hippocampal neurons ^14^. This work used a behavioral paradigm constituted of two tasks: spatial navigation and cue response. The extensive literature on HPC and DS memory systems predicts that HPC supports spatial navigation but not cue-response associations. It was therefore puzzling when a later experiment reported that damage to the medial entorhinal cortex or CA3 hippocampal field affected the ability to follow the visible cue ^15^. To systematically investigate the influence of past experience on the contributions of HPC and DS memory systems to behavior, we trained rats either individually in one of the two tasks, or concurrently in both tasks (Fig. 1A). When the animals performed to a set criterion, they received either sham surgery or selective lesions of the HPC, medial dorsal striatum (DSM) or lateral dorsal striatum (DSL). We distinguished between the medial and lateral regions of the DS since multiple data sets show that remembering cue-response associations depends mostly on DSL, while DSM supports behavioral flexibility ^16-20^ ^21^. After recovery, all animals were tested for retention across 5 consecutive days. At the end of retention testing, we further trained the animals belonging to the individual training groups in the task that would have been novel to them at that point (Fig. 1B).

**Figure 1.**
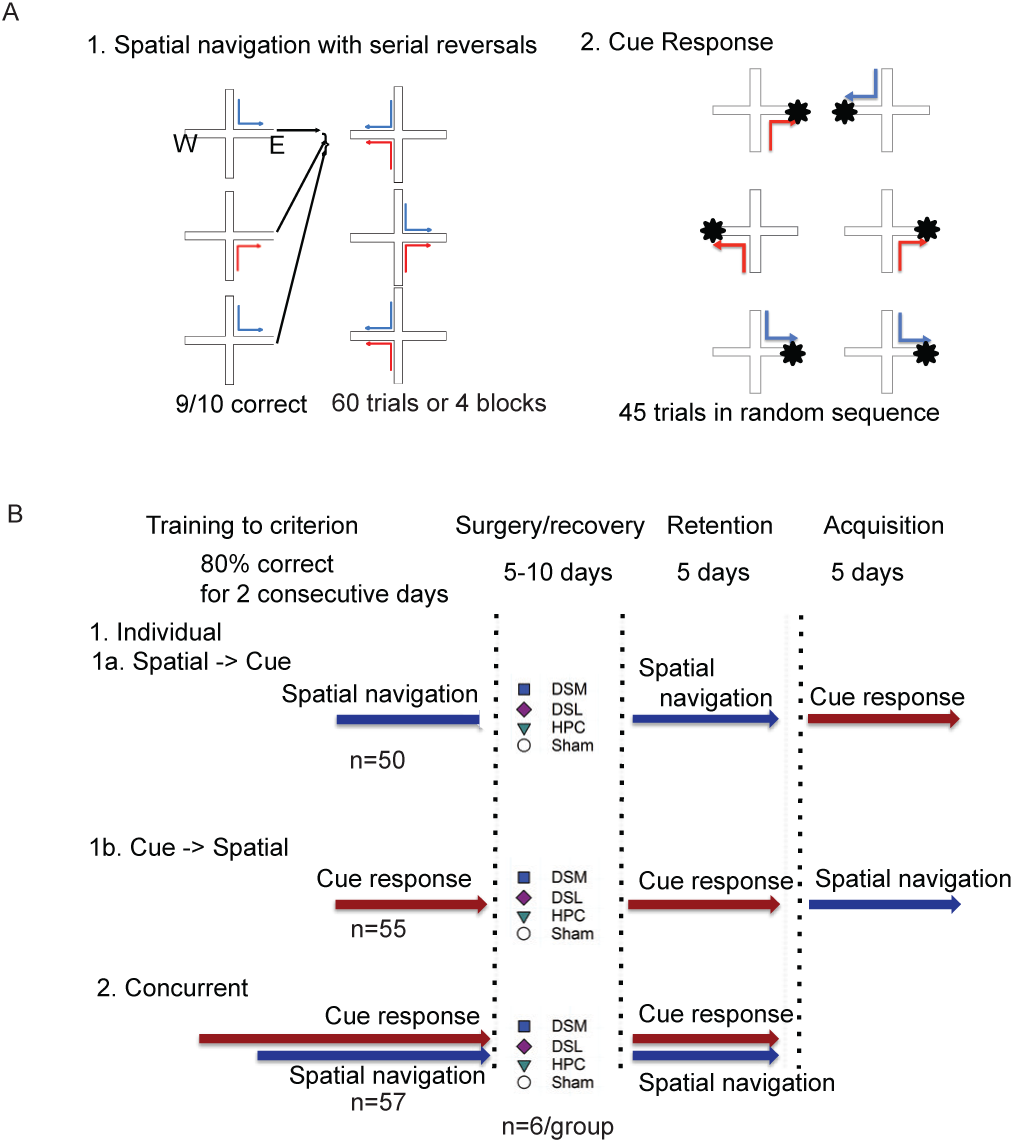
Behavioral paradigm and experimental design. **A**. The experiment used two different tasks: spatial navigation with serial reversals (1) and cue response (2). In the spatial navigation task, animals started in the North or South arm of a plus maze and walked into either the East or the West goal arms to find food based on spatial location. In the cue task, the food was placed on a white intramaze flag whose position was changed randomly from one trial to the next. **B**. Two training conditions were used, individually and concurrent. In the individual condition, rats were trained in spatial navigation task (n = 50) or cue response (n = 55). In the concurrent condition (n = 57), rats were trained in the two tasks at the same time. Post-recovery, retention was tested for 5 days successively in all animals. Animals trained in only one task were then trained for 5 consecutive days in the task new to them. After histological evaluation and exclusion of data from animals with unsatisfactory lesions, each group was constituted of six subjects.

## RESULTS

### Lesions encompassed the intended brain areas

We evaluated for each animal the lesion’s selectivity and we incorporated for analysis only data from animals with damage restricted to the intended brain areas. Cortical damage was small, if present, which was typically not the case (Supplementary Table 1). In the DS, the medial and lateral areas are not clearly delimited. Here we aimed for lesions as inclusive as possible, that would conform to known topography of anatomical connections ^22^. We considered as lesioned all striatal tissue with signs of gliosis, regardless of whether the principal neurons might have been spared at the periphery of the affected area (Fig. 2A, top two rows; Supplementary Fig. 1). As a group, the lesions distinguished well between the medial and lateral areas of the DS (Supplementary Fig. 2 A, B). Lesions of the HPC encompassed all its fields (dentate gyrus, CA3, CA1) in both dorsal and ventral areas, and most of the lesions were nearly complete. Entorhinal cortex and other nearby cortical areas were not affected (Fig. 2, bottom row; Supplementary Fig. 2C). There was minor damage to the subiculum in the anterior areas. Quantification of the lesion sizes indicated that within each lesion group, the extent of the lesions was similar across the three training conditions (DSM: F_(2,15)_ = 0.4, p = 0.6777; DSL: F_(2,15)_ = 3.26, p = 0.0666; HPC: F_(2,15)_ = 3.32, p = 0.0639; Fig. 2 inset).

**Figure 2.**
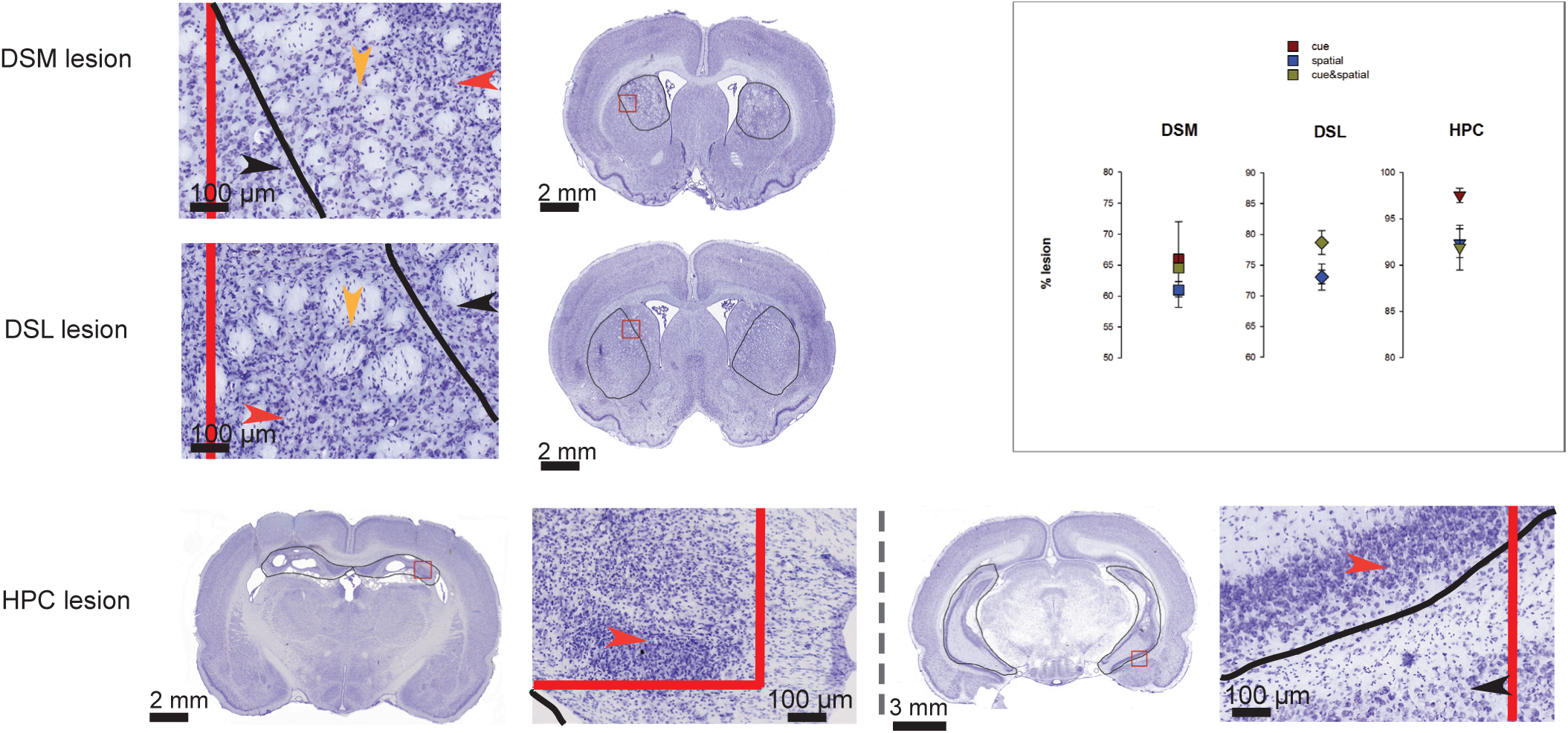
Lesions were localized in the targeted areas. Photomicrographs of coronal sections through the brain shown at low magnification indicate the lesion area (circled in black) for one example of DSM lesion (top row), DSL lesion (middle row) and HPC lesion (bottom row; left: dorsal pole, right: ventral pole). Magnified sections of the areas marked by the red squares are displayed to the left of the corresponding section. Red and black lines represent the edge of the red square and the border of the lesion, respectively, as they appear at the higher magnification. For DSM and DSL lesions, the quantification encompassed all tissue showing gliosis regardless of whether principal neurons were completely eliminated (red arrowheads) or still present among the astrocytes (orange arrowheads). Tissue considered healthy (black arrow heads) did not present astrocytic invasion. Similarly, for HPC lesions the quantification encompassed areas completely devoid of neurons (dorsal HPC, red arrowhead) and areas of neural degeneration (ventral HPC, red arrow head; compare degenerating HPC neurons to the healthy neurons in the adjacent cortex indicated by black arrowhead). More examples of lesions can be found in Supplementary Fig. 1. The results of the quantification analysis indicated that lesion size did not differ across training conditions for any experimental group (N = 6 rats/group; inset to the upper right).

### When animals were trained in individual tasks, HPC and DSL lesions selectively affected place and response memory, respectively

The memory systems theory postulates that the HPC and DSL memory systems have distinct preferred modes of information processing, which result in distinct forms of representation: S-S vs. S-R, respectively. In turn, these representations guide different types of memory-based behaviors such as spatial navigation and habits. We expected that HPC lesions would impair performance in the spatial navigation but not cue response task; and conversely, that DSL lesions would impair performance in the cue response, but not spatial navigation task. If DSM is involved in behavioral flexibility ^23^, then DSM function should be involved in both tasks to support the change in overt motor response necessary to reach the goal, as suggested by previous results ^17^ ^20^ ^24^. The empirical results obtained in the current experiment validated these predictions (Fig. 3A; Supplementary Tables 2-3).

**Figure 3.**
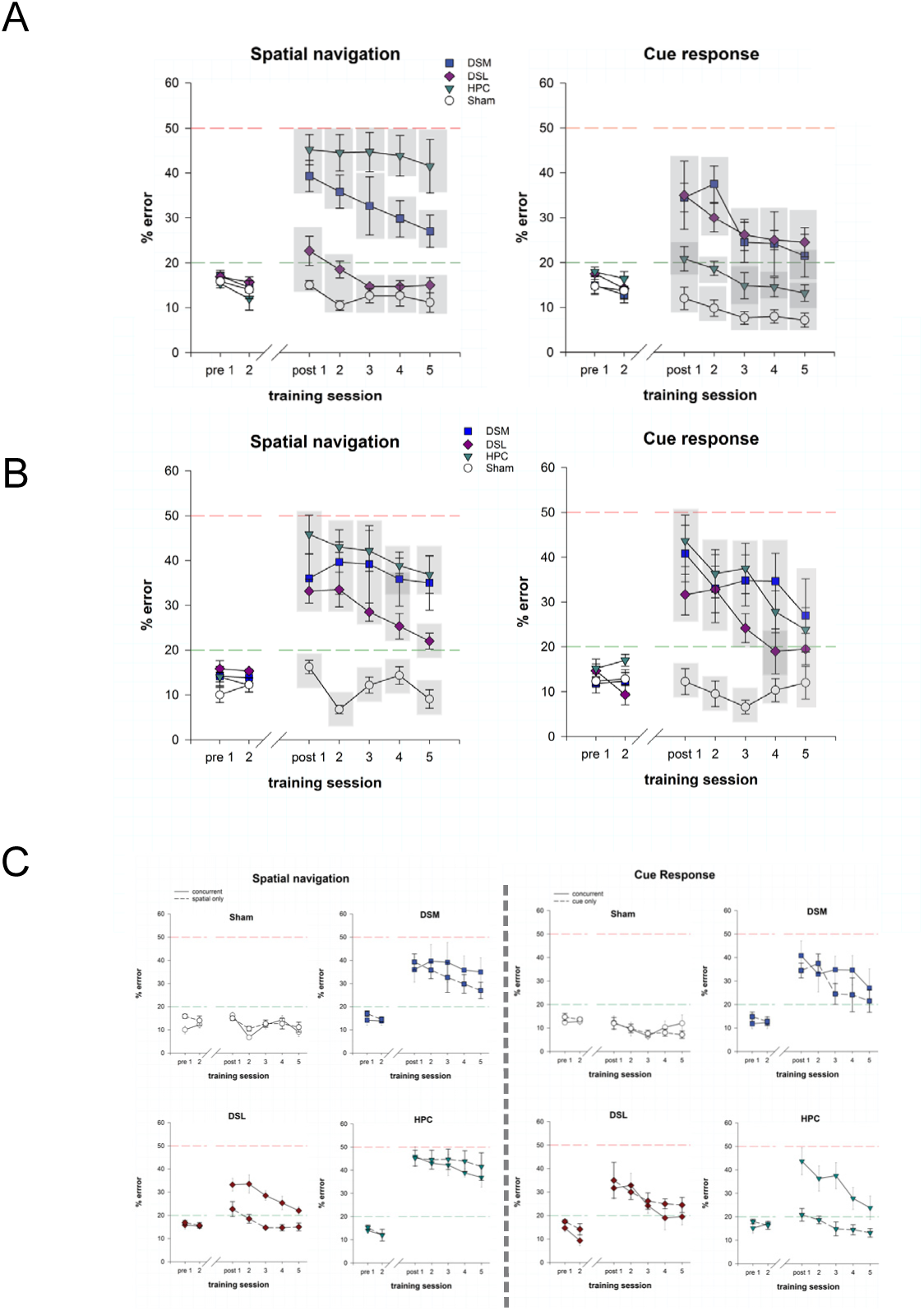
Retention in the two training conditions. **A**. When animals were trained in one of the two tasks (N=6 rats/group), HPC selectively impaired spatial memory, while DSL lesions selectively impaired cue response. DSM lesions had an effect in both cases. **B**. When animals were trained concurrently in both tasks (N=6 rats/group), HPC lesions additionally impaired cue response performance; and conversely, DSL lesions additionally affected spatial navigation. DSM continued to affect memory for both types of memories. **C**. Same data as above reorganized to facilitate comparing retention performance after individual vs. concurrent training. Vertical axis shows %error in behavioral performance (mean +/-SEM). The red horizontal line at 50% level indicates chance performance, when a rat would choose randomly between the two arms. The green horizontal line indicates the criterion threshold of 20%. Gray rectangles refer to the results of post-hoc Student-Newman-Keuls multiple comparisons test; data points included in the same rectangle are not significantly different

Overall, the lesions caused marked deficits in both cases (spatial task: F_(3,20)_= 28.92, p<0.0001; cue response task: F_(3,20)_= 12.51, p<0.0001; Supplementary Table 2), and training across days improved performance (spatial task: F_(4,80)_=4.89, p=0.0014; cue response task: F_(4,80)_=9.04, p <0.0001). In the spatial navigation task, not surprisingly, HPC lesions severely and permanently impaired performance. In contrast, animals with DSL lesions were not significantly different from controls on this task, and their performance benefited somewhat from training (F_(4, 20)_= 5.29, p= 0.0045). As expected, DSM lesions also impaired spatial navigation, initially at a level similar to the one produced by HPC lesions. The data suggested that training helped some animals to reduce the handicap, but as a group, rats with DSM lesions did not improve significantly across days and this group never attained control levels of performance. In the cue response task, DSL lesions produced a significant and persistent deficit in the cue response task. The DSM group showed relearning with extended training (F_(4,20)_=4.68, p = 0.0079) but not sufficient to reach control level. In contrast, the HPC group was not different from controls except during second day of retention testing, when it made slightly more errors (means (SE): HPC 18.67(1.62) % error vs. sham 9.83(1.80) %error); the performance of this group improved significantly during retention testing (F_(4,20)_=3.46, p = 0.0195). Thus, overall, these results validated the principle of independent parallelism proposed by the memory systems theory. The data also were in agreement with the idea that DSM is necessary for maintaining the plasticity of overt motor behavior so that it can be instrumental in reaching goals, although we found evidence of stereotyped behavior (i.e., persevering in a particular body turn regardless of circumstances) in only three of the total of twelve animals with DSM lesions.

### Concurrent training in the two tasks markedly modified the contributions of HPC and DSL to behavioral performance

If the principle of independent parallelism is universally valid, then the double dissociation pattern of results obtained when animals learn only one task at a time should continue to be present when the animals learn the tasks concurrently. However, contradicting this view, in rats concurrently trained in the two tasks described above the principle of independent parallelism was no longer valid (Fig. 3B, Supplementary Tables 1-3).

Overall, lesions had a strong effect in both tasks (spatial task: F_(3,20)_= 13.26, p<0.0001; cue response task: F_(3,20)_= 7.09, p=0.002; Supplementary Table 2) which diminished with training across days (spatial task: F_(4,80)_=5.52, p=0.0006; cue response task: F_(4,80)_=8.27, p <0.0001). HPC lesions severely and permanently impaired spatial navigation with no evidence of learning across the 5 days of retention testing, while DSL lesions interfered with performance in the cue response task. DSM lesions had as strong of an effect as HPC lesions on spatial navigation, which they permanently impaired; and had an overall marked, but somewhat less severe effect on cue response (F_(4,20)_=3.71, p = 0.0205), although three out of the six animals in this group showed strong stereotyped behavior. Thus, in these aspects, HPC, DSL and DSM continued to support behavioral performance in a pattern corroborating previous findings. However, the consistencies stopped here. DSL lesions also caused a severe spatial deficit, initially producing as much impairment as HPC and DSM lesions. The deficit diminished progressively (F_(4,20)_=4.55, p = 0.0018), but performance never reached control level. Similarly, HPC lesion impaired the cue response behavior as strongly as DSM and DSL lesions. The magnitude of the effect also diminished gradually (F_(4,20)_=3.05, p = 0.0407), but in the last day of retention testing HPC group was on average still above the 20% error criterion threshold.

We then directly compared the performance of the four experimental groups across training conditions for each of the two tasks (Fig. 3C, Supplementary Table 4). In normal animals and animals with DSM lesions, the training method had no effect regardless of the type of behavioral strategy. Overall the performance varied across days for shams (spatial task: F_(4,40)_=7.80, p < 0.0001; cue response task: F_(4,40)_= 3.01, p = 0.03) although the data did not suggest any further improvement. The DSM group did not show improvement across days in the spatial task either, but did so in the cue task (F_(4,40)_=6.47, p = 0.0004). Second, training method did not affect the spatial navigation performance of animals with HPC lesions, nor the cue response performance of animals with DSL lesions. In contrast, the training method heavily affected the performance of rats with HPC lesions in the cue response task (F_(1,10)_ =21.08, p = 0.001) and the performance of rats with DSL lesions in the spatial task (F_(1,10)_= 25.187, p = 0.0005). The magnitude of the effect diminished across days for HPC lesions in the cue response task (F_(4,40)_=5.0, p = 0.0023), but not the spatial task, while DSL lesions allowed relearning in both tasks (spatial navigation: F_(4, 40)_= 8.27, p < 0.0001; cue response: F_(4,40)_ = 5.07, p = 0.0021). Taken together, all these results, suggest that when animals learned only one task, the HPC and DS memory systems were selectively involved in guiding behavior based on S-S and S-R types of representations. In contrast, when training involved concurrent acquisition of tasks, the contributions of HPC and DS memory systems to behavior changed radically and no longer selectively supported only one type of behavioral strategy.

### Learning one type of task distinctly modulated subsequent acquisition of the other task

The results above suggest that the neurobiological basis of memory-guided behavior is not set, but depends on past experience. If so, does past experience also influence new learning? To address this issue, we shifted each of the groups in the individual training condition to training in the task that would have been novel to them at that point. The results suggest that indeed, the activity of one memory system may have long term effects on the ability of a different memory system to guide behavior (Fig. 4, Supplementary Table 5).

**Figure 4.**
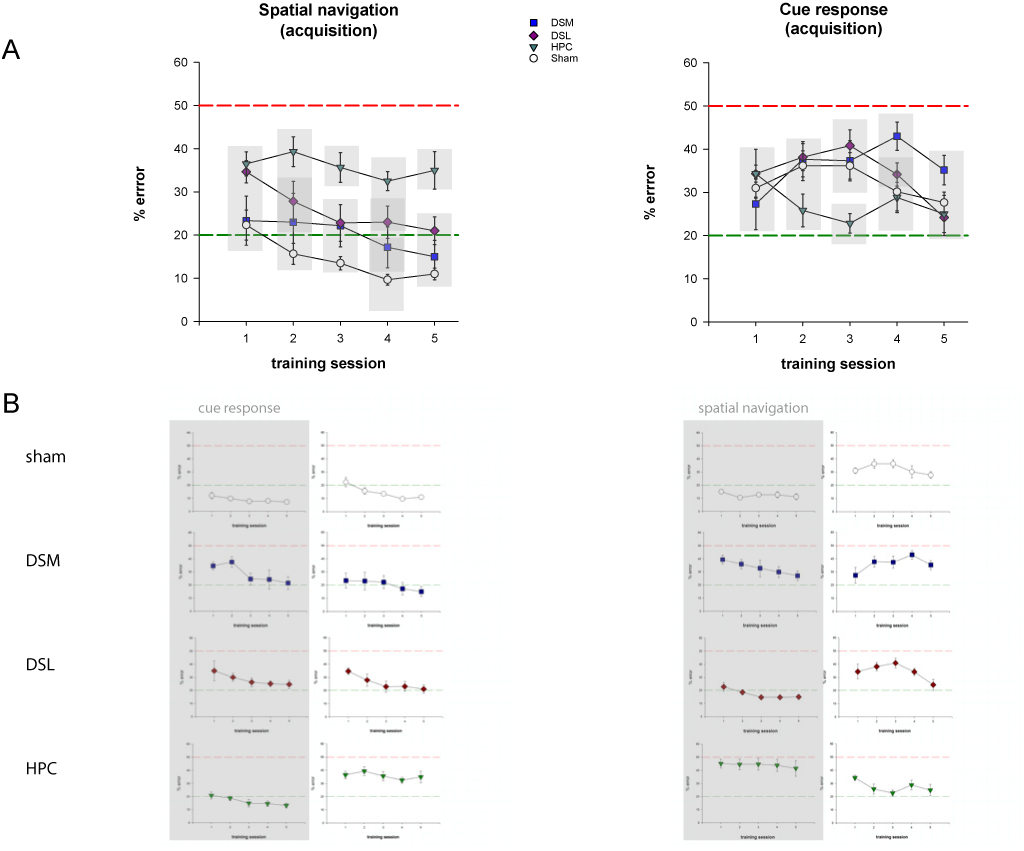
Acquisition of a new task following retention testing. **A**. After extensive experience with cue response, normal animals performed well in the spatial navigation task during the first day of learning. In contrast, experience with spatial navigation did not facilitate acquisition of the cue response task. Data are presented as in Fig. 3, N = 6 rats/group. **B**. Same data as in (A) above and Fig. 3A reorganized to emphasize the effect of previous experience (gray highlight) on acquisition of the two tasks.

The four groups differed significantly both in spatial (F_(3,20)_ = 8.73, p = 0.0007) and in cue response learning (F_(3,20)_ = 3.58, p = 0.032). Normal animals proficient at the cue response task immediately switched strategy and performed close to the 20% criterion threshold during the first day of exposure to spatial navigation, after which their accuracy further improved (F_(4,20)_=6.73, p = 0.0013). A similar process seemed to occur in animals with DSM lesions, who performed close to control levels throughout. In contrast, the DSL lesion group, who was similar to DSM group on retention of cue-response associations, could not transfer immediately the previous experience when acquiring spatial navigation, but rather gradually improved across days of repeated exposure to the task (F_(4,20)_=4.3, p = 0.0113) to eventually reach control level. The HPC lesion group, who performed at normal levels during the cue response task, was severely impaired on spatial navigation and never recovered the memory deficit. Thus, normal animals could quickly switch from a S-R to a S-S strategy based on a process independent on DSM function. Storing a cue-response association was not sufficient to help animals with HPC lesions on the spatial navigation task, on which they consistently performed poorly.

The facilitation of spatial learning after cue response training seen in the sham group may be a non-specific effect generated by familiarity with the environment and/or the basic rule of the new task (i.e., walk on the maze to collect food). If that were the case, then normal animals trained in spatial navigation should have been able to use their experience to perform well when exposed to the cue response task. However, this was not the case. Rather, the normal group well versed in spatial navigation did not perform at 20% criterion level on any of the 5 days of cue response acquisition. Among the lesion groups, none showed a statistically significant facilitating effect of past experience during the first day of cue response training, although the HPC lesion group significantly improved their performance from the first to the second day of testing (t_(12)_=2.99, p = 0.05), and during the third day, these animals performed significantly better than the other groups. This finding echoes similar effects previously reported on improved performance of S-R tasks after disruption of HPC activity ^25^ ^26^. Taken together, the results of the acquisition tests revealed a specific facilitating effect of response learning on spatial navigation which appears to be independent of DSM. In contrast, experience with spatial navigation either does not help response learning or it may even delay it, and if so, the effect may be dependent on the HPC function. To distinguish between a neutral vs. a detrimental effect, we needed to know how fast normal rats acquired the cue response task during the training phase.

### Explicit acquisition of place learning delayed S-R learning

Assessing learning during the training phase, before the surgery, included data from all animals that participated in this study (spatial navigation: n = 50; cue response: n = 55; concurrent: n = 57). At this stage, each animal was trained until achieving a performance criterion of maximum 20% error during two consecutive days. Because this procedure entailed a variable number of training session across the three groups of animals, we computed the frequency distribution of the number of days each animal needed to reach criterion (Fig. 5). Achieving criterion level in the concurrent condition required good performance in both tasks *during the same day*, which can occur at a different point in time than when the rat reaches criterion in an individual task. Therefore, for the concurrent condition, we also counted the number of days to reach 20% error or less for two consecutive days in each task, regardless of performance in the other task. The resulting frequency distributions (referred to below as ‘cue’, ‘spatial’, ‘double - cue’, and ‘double – spatial’ in correspondence to the training regimen that originated them) were compared using Kolmogorov-Smirnov tests.

**Figure 5.**
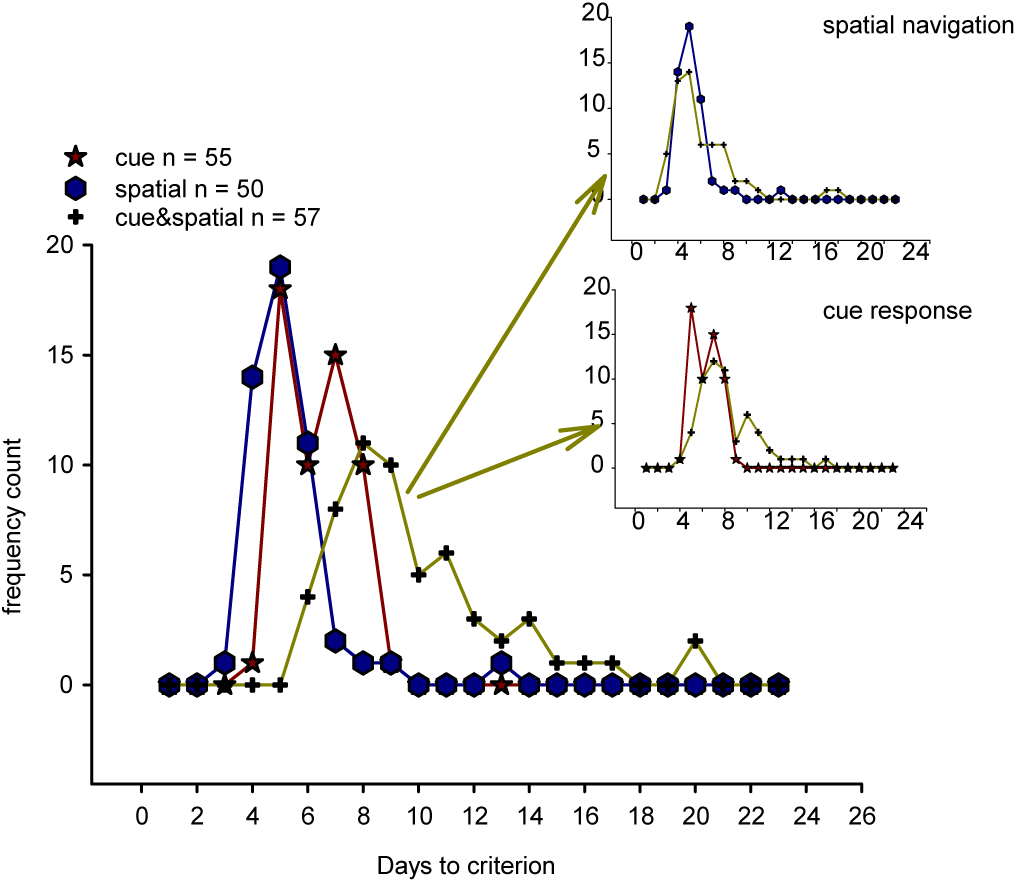
Acquisition during the initial training period. When trained in only one task, 5 days of training were sufficient for most animals in the spatial task (blue, N = 50), and a third of the animals in the cue response task (red, N = 55) to attain criterion. When trained in both tasks (dark yellow, N = 57), reaching criterion required more training, but isolating the number of days to reach criterion for each task and comparing to the corresponding number of days to criterion in the individual training condition revealed that training regimen did not affect spatial learning (upper right inset). In contrast, rats were delayed in reaching criterion on cue-response behavior when concurrently learning spatial navigation (lower right inset).

The cue frequency distribution was bimodal, indicating that some rats were ‘fast’ and some rats were ‘slow’ response learners. After 5 days of training, 33% (19/55) of the animals trained in the cue condition performed at criterion (the ‘fast’ learners, see more on this point below). In contrast, no animals (0/6) reached criterion in the same task if trained after spatial navigation (acquisition data, above). It seems therefore that previous experience with spatial learning indeed delays response learning. This result led to a second question: does *concurrent* spatial learning have a similar effect on response learning? The data suggested that this is also the case. Rats in the concurrent condition needed more time to reach criterion in the cue task than animals trained only in the same task, and after 5 days of training only 9% (5/57) of the rats in the concurrent cue condition performed at 20% or less error rate (Fig. 5, lower inset; D = 0.32, p = 0.0052).The cue and double-cue distributions were both bimodal, but the two curves were phased off by two days for the first peak and three days for the second peak. Further, if concurrent spatial learning affects response learning, is the reverse true, i.e., does concurrent cue learning have an effect on spatial learning? The data did not support this hypothesis: the spatial and double-spatial distributions were similar (Fig. 5, upper inset; D = 0.233, p = 0.11). Moreover, these curves were unimodal, indicating that there are no ‘slow’ and ‘fast’ spatial learners. How does then the rate of learning the spatial task compare to the ‘fast’ and ‘slow’ rate in the response task? The cue and spatial distributions (Fig. 5, main) were, unsurprisingly, widely dissimilar (D = 0.3727, p = 0.0015). However, both medians occurred at the 5 days time point, which corresponds to the the ‘slow’ response category. At the 5 day mark, 38% (19/50) of the rats in the spatial group and 33% (19/55) of the rats in the cue group had acquired the task. On average though, response learning developed slower than place learning (means (stdev): 6.33 +/-1.21 days vs. 5.28 +/- 1.58 days, respectively) because 30% (15/50) rats needed more than 5 days to learn the spatial task, while the corresponding number for the cue response task was 65% (36/55). Thus, the results of this analysis suggest that exposure to spatial learning, prior or concurrent, delays S-R learning while the vice-versa is not true; S-R learning occurs ‘slow’ or ‘fast’; and ‘fast’ S-R learning rate is similar to spatial learning rate.

## DISCUSSION

The present study used similar procedures as the ones previously used to generate the large body of data leading to the memory systems theory (e.g., neurotoxic lesions). Thus, our results not only integrate completely within that literature, but also are based on complete and permanent disruption of neural function in large brain areas such as the HPC and DS, a condition desirable for evaluating both the contribution of such structures to memory-guided behavior ^27^, and the function of the spared circuitry, with its compensatory abilities. During any given experience, multiple memory traces are formed and stored through the parallel activity of the memory systems; depending on circumstances, one of these representations will control behavioral output ^28^. The process of selection, crucial for the memory circuits to accomplish their function, remains however poorly understood. HPC and DS memory systems can interact either competitively or synergistically to direct behavior ^29^, ^30^ ^25^ ^28^ ^31^ ^32^ ^33^. Multiple factors are involved in these interactions ^34^ ^35^, and the current state of the field is in some aspects similar to the state of hippocampal research in the early ‘70s, before the publication of O’Keefe and Nadel’s theory of hippocampus as a cognitive map ^12^: we have many pieces of the puzzle, but we do not yet know how they fit together. A recently formulated new model of interaction between memory systems may turn out to provide the organizing theoretical framework. Starting from a putative two-process model for HPC contribution to context conditioning ^36^ and based on empirical results suggesting information transfer between memory systems ^37,38^, White and collaborators elaborated on their earlier work to propose an expanded parallel processing model in which the HPC, DS and amygdala memory systems, while still acting independently and in parallel, can also interact indirectly, through the common cortical representation all systems contribute to ^35^ ^28^. Our current data lend support to this theory. When animals learn one task at a time, operational parallelism is predominant because there is only one cortical representation formed. When animals learn two tasks concurrently in the same environment, the memory systems interact co-operatively and in synergism because of the two similar (though not identical) cortical representations, which can ‘pull’ the memory system with incongruent type of activity in the neural circuit supporting behavior (Fig. 6). Second, the model can also explain the protracted interaction between DS and HPC memory systems we currently found. The cortical representation formed when the animal learns one of the tasks can influence novel learning as it is reactivated by the identical environment within which this novel learning occurs.

**Figure 6.**
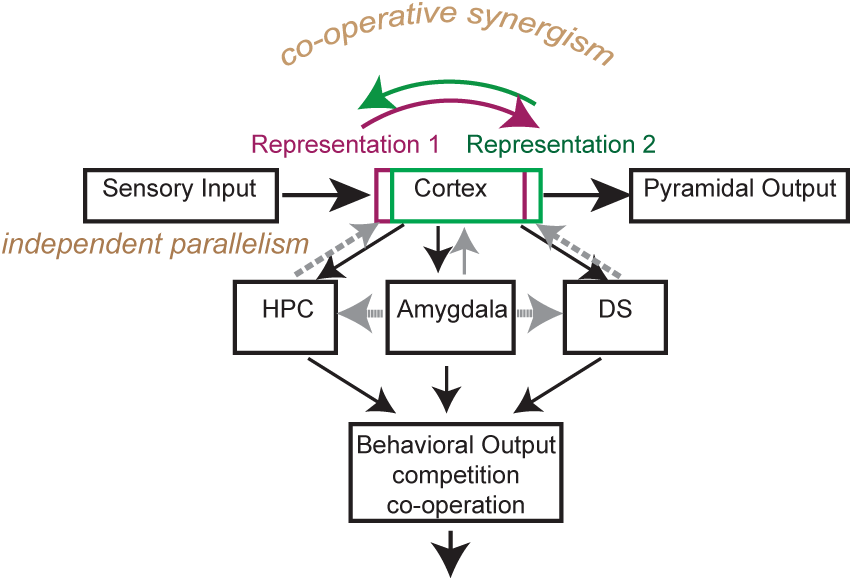
Expanded parallel processing model applied to two similar mnemonic representations. The three memory systems (HPC, amygdala, and DS) receive information from the cortex, process it in parallel, and use it to guide behavior (black arrows). Amygdala also modulates the HPC and DS memory systems (gray horizontal arrows). The model proposes that output from all memory systems is sent back to the cortex and is incorporated in the representation of a given experience. Through the cortex, memory systems can indirectly influence each other. Here, the green and the magenta overlapping rectangles represent two similar (but not identical) experiences, such are likely the ones formed during the spatial navigation and cue-response tasks in the current paradigm. For such similar experiences, feedback from the memory systems to the cortex is received by neural circuits sharing large part of their individual neurons. Consequently, activity of the HPC memory system may become involved in the performance of the cue-response task and vice-versa, activity of the DS memory system may become involved in spatial navigation, generating co-operative synergism. Adapted from White (2004).

DSL, also known as the sensorimotor striatum because of its connections with the sensorimotor cortices, has been primarily linked to memory for S-R associations and habits, a model-free type of behavior based on automatism. In contrast, DSM has been associated with model-based behavior, which is responsive to outcome and which involves behavioral flexibility ^16^ ^21^ ^19^ ^39^ ^18^ ^17^ ^20^. Several physiological recording experiments also linked DSM activity and choice ^40^ ^41^ ^42^ ^43^. The current results support this view. We found that DSM was involved in both spatial navigation and cue response regardless of training experience, a finding compatible with a role for DSM in supporting appropriate changes of body turns in pursuit of the goal. However, systematic stereotyped behavior (i.e., perseverance of a specific body turn) caused poor performance only in a relatively restricted subset of the animals with DSM lesions; in some cases, the rats relinquished the body turn strategy with training. Furthermore, DSM function did not seem to be required for normal animals to switch from cue response to spatial navigation. Thus, DSM seems to be working along other neural circuits to mediate behavioral flexibility.

According to the current view, habits interfere with, and eventually supersede behaviors based on cognitive type of memory. In contrast, our data suggest that habits may actively contribute to model-based behavior in certain conditions. Normal rats and rats with DSM lesions extensively trained in the cue response task were able to immediately adapt when they were exposed to a spatial task, while lesions of the habit-supporting DSL interfered with this ability. On the other hand, experience with a cognitive strategy consistently impeded S-R learning, raising the possibility that interference from the HPC memory system may be part of the explanation behind the slowness of habit formation processes. The bimodal nature of the cue and double-cue distributions, and the shift of the ‘fast’ response learning to ‘slow’ rate after experience with the spatial task raises this intriguing possibility.

## >METHODS

### Subjects

Male Long-Evans rats (300-350g, 4-6 months old, Charles River and Harlan) were housed in individual cages (12 hour light cycle; animals were tested during the day cycle). After acclimation, the animals were food deprived to 85-90% of *ad libitum* body weight and maintained at this level for the rest of the procedures (allowing adjustment for normal growth). All procedures met NIH guidelines and were approved by the Institutional Animal Care and Use Committee. A total of 164 animals participated in this study, but memory retention was evaluated in 18 sham controls and 54 rats with brain selective and inclusive lesions (see below) of the targeted areas (6 subjects/lesion group/condition). Acquisition of the behavioral paradigm, which occurred before the surgery, was evaluated based on the performance of the entire group of 164 rats.

### Apparatus

The plus maze was made of gray polyvinyl chloride (PVC) and elevated 91 cm from the floor of a room that contained several visual cues. Each of the 4 arms was 61 cm long and 6.3 cm wide. A grey PVC block (30.4 cm high, 6.3 cm wide, 15.2 cm deep) was used to block the start arm that was not in use for that trial. A rectangular waiting platform (32cm x 42cm) was placed next to the maze. In the cued version of the task, a white visible flag also made of PVC was used to indicate the location of the food on the maze; during the spatial version of the task, the cue was placed on a table in the room and off the maze.

### Behavioral Training and Testing

All animals were pre-exposed to the maze in the presence of food for two consecutive days and then trained to walk from either the North or the South start arms to the end of West or East goal arms to obtain half a Fruit Loop. Between trials, the rats were placed on a side platform to wait for the next trial. Entry with all four paws into the unrewarded arm defined an *error*, which the rat was allowed to correct. Training procedures followed previously published protocols ^44^ ^14,15^ and utilized a spatial navigation task with serial reversals and a cue-response task (Fig. 1). In both cases, the start and the goal arm were selected based on a pseudorandom sequence of 60 trials with ≤ 3 consecutive repetitions of the same type of journey (NE, NW, SE or SW). *Spatial task*. In the spatial task, the animal was rewarded for remembering spatial location. The position of the food was kept constant until the rat entered the correct goal arm in 9 of 10 consecutive trials. At that point, the other goal arm was baited and a new block of trials began. If the animal did not reach the criterion in maximum 20 trials at the beginning of training, or 15 trials once he was familiar with the task, the location of the food was changed automatically, to avoid unbalanced reinforcement of any specific goal arm. Alternating trial blocks continued up to either 4 blocks or 60 total trials. *Cue task*. In this case, the rats had to remember an association between the visible cue, whose location was rendered irrelevant by changing start and goal positions based on a pseudorandom sequence, and a motor response which was the walk towards the cue. Animals received 45 trials in each session, a number approximately equal to the number of trials necessary to complete four blocks of trials in the spatial task. Thus, each goal arm was rewarded approximately equally both within and across tasks.

***Experiment 1: Retention in spatial navigation task; acquisition of cue response***. One set of animals was first trained in spatial navigation only. Training continued until the rats reached a criterion of 80% correct choices for two consecutive days, after which they were assigned to one of four groups: HPC lesions, DSM lesions, DSL lesions, and sham controls. After a recovery interval of 5-10 days, retention of the spatial task was evaluated for 5 consecutive days and then all animals were trained in the cue response task.***Experiment 2: Retention in cue response task; acquisition of spatial navigation***. Procedures were similar as in *Experiment 1*, except that animals were first trained and tested in the cue response task, and then acquired spatial navigation. ***Experiment 3: Concomitant training in spatial navigation and cue response tasks***. All rats started training in the cue task. When the animals readily responded to the cue (two to three days), they were introduced to the spatial navigation task and continued with daily training in both tasks. The order in which the two tasks were presented within a session was varied to avoid an order effect. When the rats reached 80% correct performance in both tasks for two consecutive days, they were assigned to one of four groups: HPC lesions, DSM lesions, DSL lesions, and sham controls. Post-recovery, retention was evaluated for 5 consecutive days in both tasks.

### Lesions

Rats were anesthetized with isoflurane and diazepam or midazolam (10mg/kg). Atropine (5 mg/kg body weight) was also administered in order to avoid fluid accumulation in the respiratory tract. Neurotoxic lesions were made by injecting either a solution of 5mg/ml NMDA in phosphate buffer (pH 7.4); or a quinolinic acid (30mg/ml in phosphate buffer titrated with sodium hydroxide to pH 7.4) through a 30-gauge cannula attached to a minipump (0.2 μl/min; New Era Pump Systems, Inc., Model NE-4000). At the end of each injection, the cannula was left in place for 3 mins, retracted 0.5 mm and left in this location for 1 min, and then slowly retracted completely. The coordinates of each injection and the volumes injected are presented in Table 1. In order to prevent seizure development, a second, ip., injection of valium (10 mg/kg body weight) was administered prior to neurotoxin infusion and animals were monitored until completely awake and active in their home cages. Sham animals were anesthetized, incised and sutured.

**Table 1.**
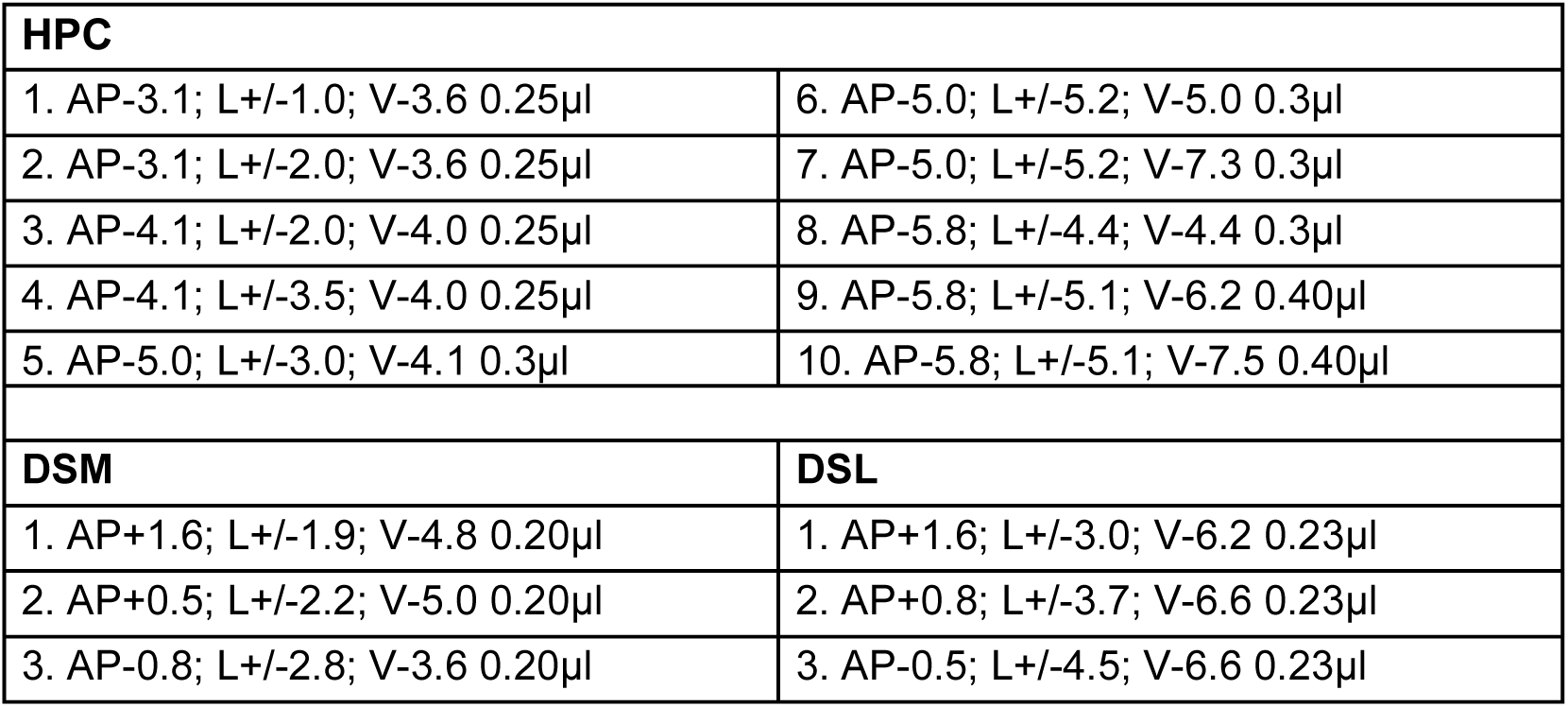
Lesion coordinates. AP, antero-posterior from bregma; L, lateral from bregma; V, ventral from bregma. All coordinates are in mm.

### Behavioral Data analysis

Percent performance error was calculated for each rat during each day of testing and a mean was calculated for each group during each day. Differences in performance were assessed using two-way ANOVA analyses (The SAS Institute). Time (in days) and lesion groups as categorical independent variables; and performance as percent error on repeated time points as a dependent variable were entered into the model. Where overall analyses indicated significant differences, we focused on simple effects; where simple effects were significant, we proceeded to performed multiple comparisons adjusted by the Student-Newman-Keuls procedure. We also investigated learning during the training phase (before surgery) by computing the frequency distributions for number of training days to criterion in each of the three training conditions. We compared these distributions using a Kolmogorov-Smirnov test (The SAS Institute).

### Tissue Preparation and Processing for Histology

Rats were overdosed with isoflurane (administered in closed environment) and perfused transcardially with normal saline, then 10% formalin for tissue fixation. Coronal sections (40 μm) were cut on a cryostat and stained with cresyl violet to evaluate the extent of the lesion.

### Lesion Assessment

Brains were sectioned coronally. Each 4^th^ section in the striatum lesion groups, and each 5^th^ section in the hippocampal lesion groups was mounted on a microscope slide to be used for lesion evaluation. We first inspected visually under the microscope and traced each lesion on a set of histological plates ^45^. Data from animals whose lesions were not sufficiently inclusive (i.e., encompass most of the targeted area) and selective (i.e., extended bilaterally to significant portions of other brain areas) were excluded from further analysis. The lesions that passed criterion underwent quantification analysis. We used for this purpose every second preserved section from rats with hippocampal lesions and every third preserved section from rats with striatal lesions. For each of these brain sections, we overlaid a 17x34 grid on a 20X magnification image of the area of interest (600X800 pixels) and counted the grid units that covered healthy tissue. This procedure was also performed for brain tissue from two control animals for the dorsal striatum and two control animals for the hippocampus. The number of grid units from the control animals were summed up to obtain the total DS or HPC areas, respectively, and then the two values were averaged to obtain the value estimating healthy HPC and DS ‘volumes’, respectively (because the brain tissue was sampled at regular intervals of 200 μm for all HPC tissue and 160 μm for all DS tissue in control and lesion animals alike, multiplying by the constant distance between sections was not necessary to obtain an estimate of the lesion as % from the total structure). %healthy tissue was computed by dividing the number of grid units covering healthy tissue in a lesioned brain by the number of grid units corresponding to the area in a control brain; the %lesion was obtained then as the difference from %healthy tissue to 100%. For each lesion groups we compared the %damage across the three experimental conditions using one-way ANOVA (The SAS Institute).

## ACKNOWLEDGEMENTS

We thank Yitzhok Becher, Judah Horowitz, Maiko Iijima, Khiara Scolari, Metika Ngbokoli, and Isaac Buff for help with animal training; and Fraser Sparks for help with lesion procedures. This work has been supported by NIH grants MH106708 and MH094946, and by an internal SUNY Downstate grant.

